# Coincidence of cholinergic pauses, dopaminergic activation and depolarization drives synaptic plasticity in the striatum

**DOI:** 10.1101/803536

**Authors:** Yan-Feng Zhang, Simon D. Fisher, Manfred Oswald, Jeffery R. Wickens, John N. J. Reynolds

## Abstract

Pauses in the firing of tonically-active cholinergic interneurons (ChIs) in the striatum coincide with phasic activation of dopamine neurons during reinforcement learning. However, how this pause influences cellular substrates of learning is unclear. Using two *in vivo* paradigms, we report that long-term potentiation (LTP) at corticostriatal synapses with spiny projection neurons (SPNs) is dependent on the temporal coincidence of ChI pause and dopamine phasic activation, critically accompanied by SPN depolarization.

---

The pause response in striatal tonically active neurons, believed to represent cholinergic interneurons (ChIs), coincides with phasic activation of midbrain dopamine neurons during learning^1,2^. Phasic dopamine activity plays a critical role in plasticity at synapses between cortex and striatum *in vivo*^3,4^. *However, how the pause response in ChIs interacts with phasic dopamine to induce corticostriatal synaptic plasticity in vivo* remains unclear.

Elucidating the functional significance of pauses has been hindered by the challenge of inducing synchronized pauses in sparsely-distributed ChIs, aligned with phasic activity of dopamine neurons. We recently showed that the firing of ChIs is entrained to fluctuations in excitatory input^5^, with the inverted striatal local field potential (iLFP) providing a read-out of spontaneous and electrically-evoked excitatory activity in our anesthetized preparation. Here, using the iLFP as a read-out of firing patterns of ChIs, we investigated the effect of phasic dopamine occurring at different phases of the ChI pause response.

Firstly, we made single unit extracellular recordings of SPNs *in vivo* (Figure 1A, B & D) and measured corticostriatal plasticity by observing changes in the probability of eliciting a cortically-evoked spike. Cortical electrical stimulation applied under urethane anesthesia also entrained the firing of striatal ChIs, with the pause indicated by the ‘receding phase’, and maximal firing during the ‘rising phase’, of the iLFP^5,6^ (Figure 1C). To drive phasic activity in dopamine neurons, we applied a light flash as a salient visual event to the contralateral eye, with the superior colliculus (SC) disinhibited by local injection of the GABA antagonist bicuculline (BIC; Figure 1A). During collicular disinhibition, visual stimuli drive phasic activity in dopamine neurons, about 110 ms after a light flash^7,8^. During the pairing of cortical and visual stimulation, in order to align phasic dopamine activation to a pause (Match group), or excitatory phase of ChIs (Mismatch group; Figure 1C), we manipulated the interval between the stimulations so that the light-driven dopamine phasic activity occurred during the ‘receding’ or ‘rising’ phase of the iLFP, respectively.

**Figure 1.**
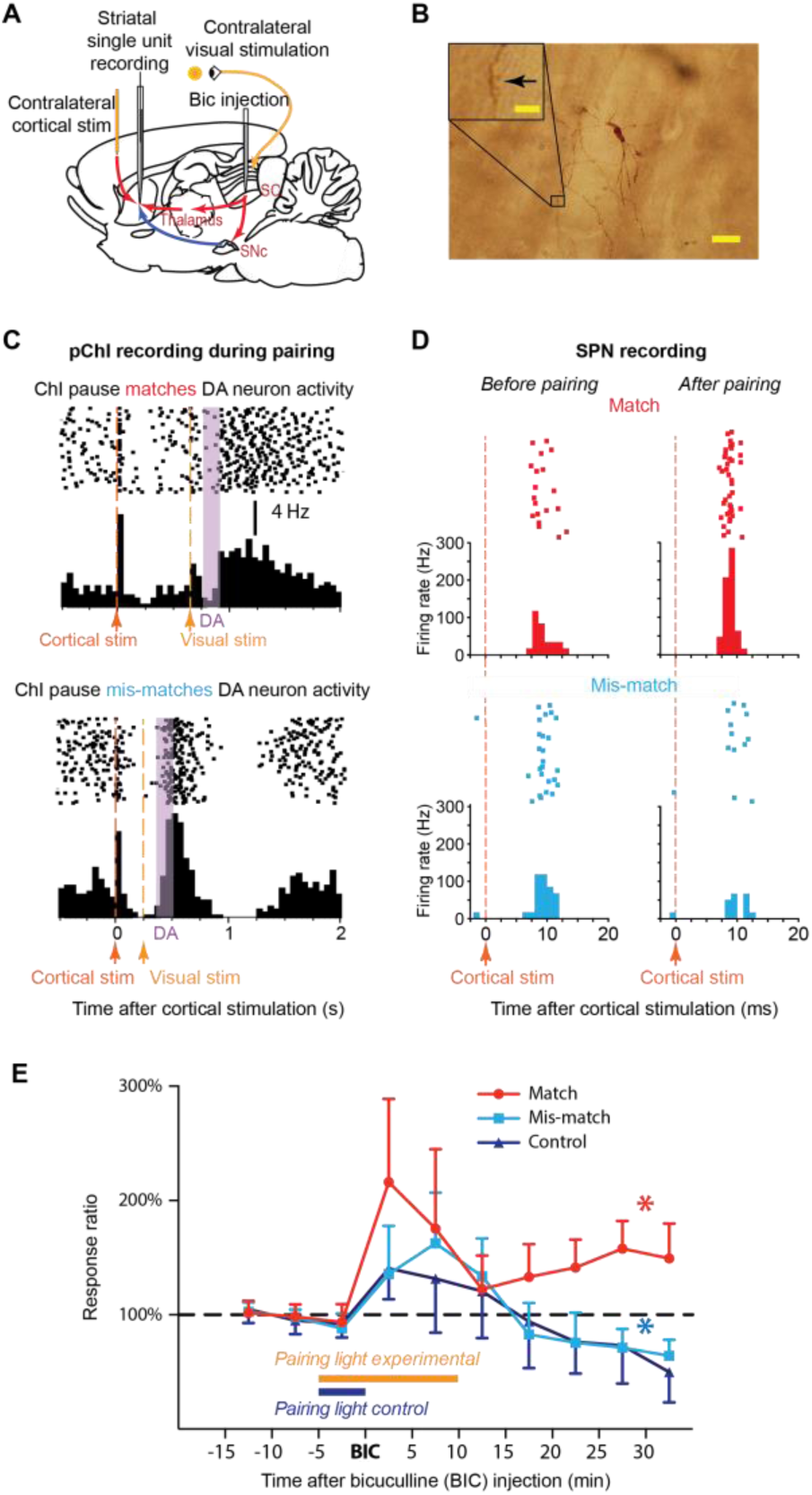
SPN spikes induced by cortical stimulation were potentiated by coincident ChI pause, dopamine activation and depolarization. (A) Cortical stimulation induced spike activity in SPNs. Contralateral visual stimulation with BIC injected into the SC activates dopamine neurons and depolarizes SPNs. (B) Example labelled SPN with spine arrowed in inset (Scale bar 20 µm; inset 5 µm). (C) Peristimulus time histograms (PSTH) of ChI firing activity during stimulus pairing. The estimated time of dopamine phasic activity was aligned to ‘match’ (upper) or ‘mis-match’ (lower) the ChI pause. (D) A representative potentiation (upper) and depression (lower) in SPNs 30 minutes after match or mis-match pairings, respectively. (E) Spike activity in SPNs was potentiated when the light-induced dopamine signal was coincident with ChI pauses (red), but not when coincident with excitation of ChIs (light blue) or when phasic dopamine was absent (dark blue - no light flashes after BIC). *p<0.05 (paired t-test compared to baseline).

With ChI pauses and phasic dopamine coincident in the ‘Match’ group, potentiation of cortically-induced SPN spike activity was induced (Figure 1D upper & 1E,). In contrast, depression of spike activity occurred in the ‘Mismatch’ group (Figure 1D lower &1E), in which dopamine was temporally separated from the ChI pause. In the control group (‘Control’), light flashes were only paired with cortical stimulation for 5 min prior to bicuculline injection, so the dopamine neurons were not phasically-activated during the pairing protocol. Thus, ChI pauses elicited by cortical stimulation were insufficient to induce potentiation without phasic dopamine, instead inducing a similar depression to the ‘Mismatch’ group (Figure 1E). Therefore, potentiation of corticostriatal responses only emerged when phasic dopamine was coincident with the ChI pause.

Notably, a light flash applied during collicular disinhibition also depolarizes SPNs via the tecto-thalamo-striatal glutamatergic pathway (Figure 1A)^9,10^. Thus, this depolarization would occur at the same time as the dopamine signal. To disentangle the effects of SPN depolarization and dopamine on corticostriatal plasticity, in place of collicular disinhibition we used intracellular current injection to manipulate SPN depolarization independently of SNc dopamine cell activation (Figure 2A & B).

**Figure 2.**
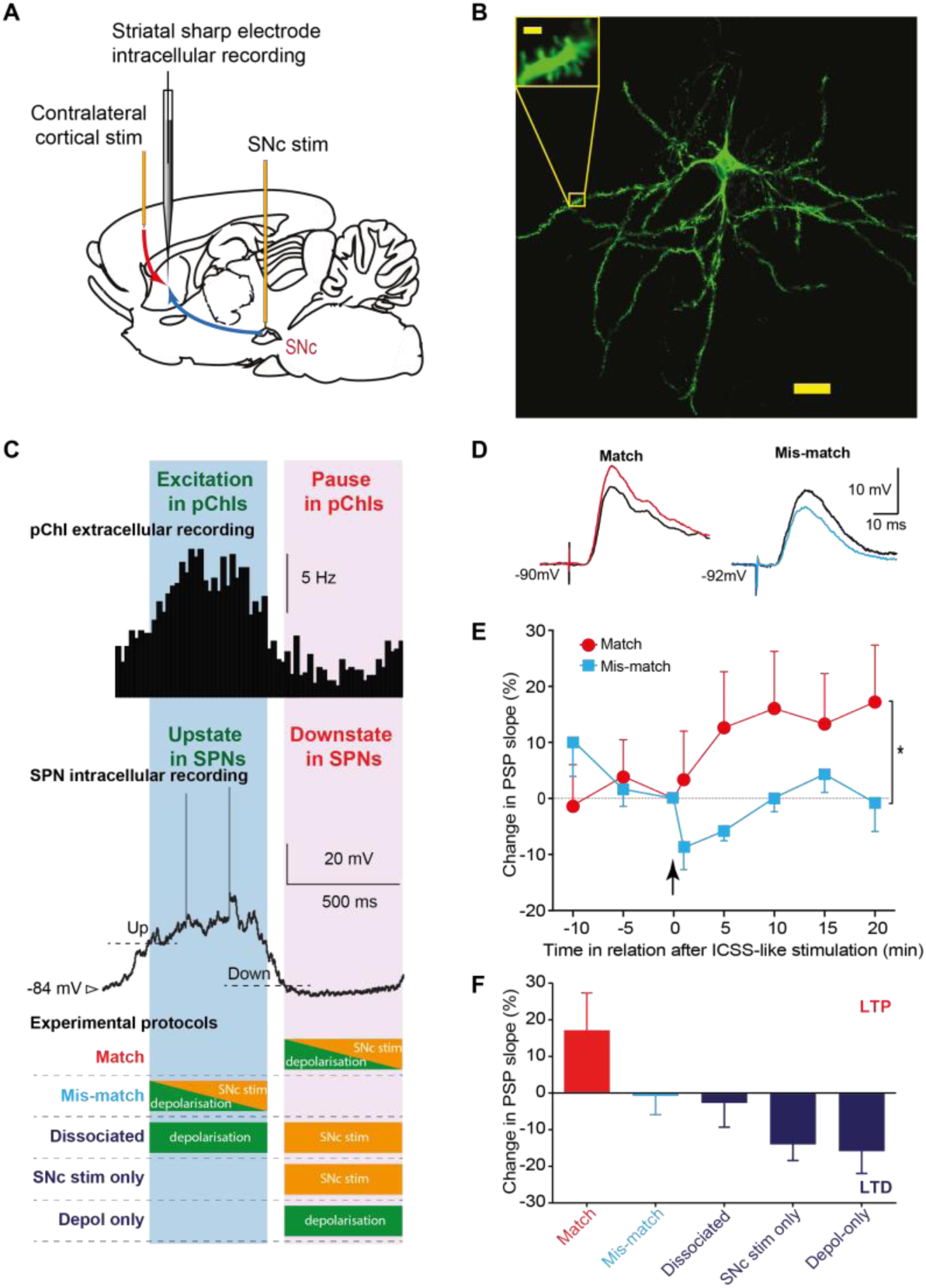
Coincidence of ChI pause, dopamine activation and depolarization potentiated corticostriatal SPN postsynaptic potentials (PSPs) (A) Cortical stimulation induced PSPs in SPNs. Dopamine neurons were activated by electrical stimulation and depolarization was induced by intracellular current injection. (B) Example labelled SPN (Scale bar 20 µm; inset 2 µm). (C) Electrical stimulation of the SNc was set to match (purple shade) or mis-match (blue shade) the ChI pause, determined from the striatal iLFP. (D) Example corticostriatal PSPs potentiated (red) or depressed (light blue) 20 minutes after baseline (black traces) when SNc stimulation matched or mismatched the ChI pause, respectively. (E, F) Mismatched SNc stimulation with ChI pause, dissociated SPN depolarization and SNc stimulation, SNc stimulation alone, and depolarization alone all induced depression. *p<0.05 (repeated measures ANOVA; N=5 to 7 neurons per group; post hoc comparisons between groups p<0.01).

Previously, we demonstrated an association between synaptic plasticity and positive reinforcement^4^. In that study, dopamine neurons were activated simultaneously with SPN depolarization during the SPN ‘down state’, which corresponds to a period of minimum excitatory cortical input to the striatum^11^. We recently demonstrated it is also the time when ChIs exhibit a pause^5^ (Figure 2C).

Here, we investigated the requirements for synaptic plasticity further using the same paradigm. Consistent with our previous work^4^, LTP was induced by combined dopamine cell stimulation and depolarization during a period of relative corticostriatal silence (‘Match’: 17.2 ± 10.2% at 20 min, n=7 rats; Figure 2C, D & E). In contrast, no significant change in synaptic efficacy resulted when SPN depolarization and dopamine input were applied during the spontaneous SPN up state, at the time when ChIs are most excited (‘Mismatch’: −1.0 ± 5.1% at 20 min, n=7; Figure 2E). These results agree with the extracellular experiments in demonstrating a need for dopamine activation to coincide with the pause in ChIs to induce lasting potentiation.

We then investigated if coincidence of dopamine cell stimulation, ChI pause and depolarization of SPNs is necessary to induce LTP at corticostriatal synapses. With either dopamine stimulation alone (SNc stim only) or depolarization of SPNs alone (Depol only) applied during the ChI pause period, LTD resulted. Further consistent with the need for temporal coincidence, depolarization of SPNs by current injection during corticostriatal excitation, when combined with SNc stimulation during the pause phase of ChIs (’Dissociated’), induced no change in synaptic efficacy Figure 2C & F).

In summary, we investigated in intact animals the influence of synaptically-driven ChI pauses on corticostriatal synaptic changes relevant to task learning. By examining ChI pauses entrained by excitatory input rather than induced by optogenetic manipulation^12-14^, we were able to characterise the effect of pause responses naturally flanked by endogenous excitations, which may themselves encode reward-relevant information^15,16^. In addition, visual stimulation in experiment 1 enabled us to trigger dopamine activity^8^ and concomitant thalamostriatal depolarization^9^ via physiological pathways. Intracellular recordings in experiment 2 further allowed us to dissociate the effects of each input during plasticity induction.

Despite differences in recording techniques and measures of plasticity, results from both experimental preparations were in agreement that a temporal coincidence of phasic dopamine, ChI pause and SPN depolarization is required for the induction of corticostriatal LTP *in vivo*. Notably, the essential depolarization accompanying the dopamine input arrived after the corticostriatal-driven “Up” state. This is consistent with our recent report that corticostriatal potentiation is optimally induced by a sensory stimulus arriving within a critical temporal window following presynaptic corticostriatal activity and SPN firing^17^. The present study demonstrates how the ChI pause shapes these critical temporal constraints (Figure 3).

**Figure 3.**
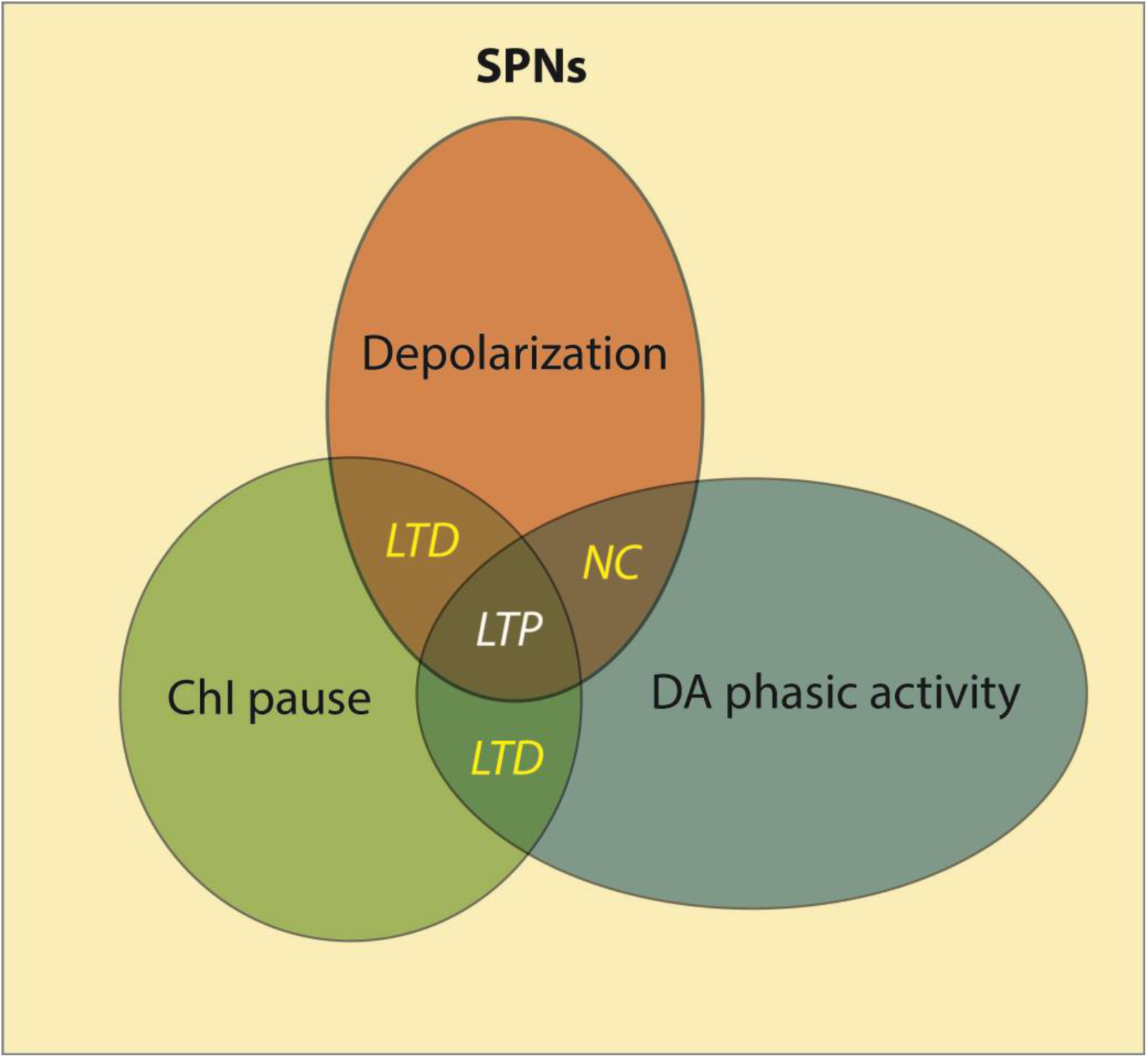
A model of corticostriatal synaptic plasticity. Corticostriatal LTP is induced when SPNs are depolarized (brown) in a striatal area where a ChI pause response (green) is coincident with a phasic dopamine signal (blue). Other conditions will result in LTD or no change (NC).

Phasic dopamine activity, believed to report prediction error^18^, is widely broadcast throughout the striatum by rich striatal arborizations. ChIs are sparsely distributed in the striatum and develop a synchronized pause response to discrete excitatory stimuli in a subset of the striatal territory^19^, which can be enhanced by overtraining of reward associations^20^. Our data suggest that SPN synaptic potentiation may be directed by the pause response of the ChIs to the striatal location where new stimulus-reward associations have become established during learning, through the effect of a localised dip in acetylcholine tone. In contrast, other synapses will be unchanged if located outside the area of the pause or depressed if phasic dopamine is absent or arrives without an accompanying sensory depolarization (Figure 3). In summary, our study provides the first empirical data supporting the important temporal gating of corticostriatal synaptic plasticity by a physiologically-induced pause response in ChIs.

## Acknowledgements

This work was supported by grants from the Marsden Fund of the Royal Society of NZ (to JNJR, and JRW/JNJR), Lottery Health Research (JNJR/JRW) and the Neurological Foundation of New Zealand (MJO/JNJR). Our thanks to Annabel Kean and Koreen Clements for technical assistance, and to Jan Schulz, Peter Redgrave and Gordon Arbuthnott for helpful discussions that have shaped this manuscript.

## Methods

All procedures in this study were conducted in accordance with approvals granted by the University of Otago Animal Ethics Committee. A total of 136 male Long-Evan rats for extracellular recording were used, yielding 258 putative spiny projection neurons, and 18 were successfully recorded for the full plasticity recording protocol. Intracellular recording was performed with 105 male Wistar rats, yielding 32 spiny projection neurons recorded for the plasticity protocol.

### Surgery

Male Long–Evans rats (250–450 g) or Wistar rats (280 – 450g) were anaesthetized with urethane (1.4-1.9 g/kg i.p.; Biolab Ltd., Auckland, New Zealand). During recording, the level of anaesthesia was monitored by continuous observation of the band-pass filtered EEG signal (0.01 to 500 Hz). Supplementary urethane was administered via an intraperitoneal catheter at any sign of EEG desynchronization, indicating a lessening of depth of anaesthesia. The head was fixed in a stereotaxic frame (Narishige, Japan) and core temperature maintained above 36 °C by a homoeothermic blanket and rectal probe (TR-100, Fine Science Tools). All wounds and pressure points were infiltrated with a long-acting local anaesthetic (Bupivacaine, 0.5%).

For monitoring of the electroencephalogram (EEG), a hole was drilled in the skull above the left posterior cortex, and a silver wire electrode placed against the dura overlying the cortex and fixed in placed with dental cement. A flap of bone overlying the cortex was removed to provide access to the recording site in the left medial striatum, and a “well” of dental cement fashioned around the perimeter of the hole. All coordinates are given in millimetres in relation to Bregma and the midline.

### Electrical stimulation

To implant a stimulating electrode into the medial agranular motor cortex, a round piece of skull overlying the right hemisphere (centred AP +2.0 to +2.7 mm and ML −1.6 to −2.0 mm to Bregma) was removed. A concentric (extracellular experiments; Rhodes NEW-100X 10 mm, USA) or parallel-contact (intracellular experiments, locally manufactured) stimulating electrode was implanted in the medial agranular motor cortex to a depth of 1.6 to 2.4 mm. Stimulating electrodes were connected to constant current electrical stimulators (Isolator-10, Axon Instruments Inc.) Stimulus pulses applied to the cortex were biphasic (0.1-0.2 Hz, 0.1 ms, 300 to 990µA). For experiments requiring substantia nigra stimulation, the medial contact of a parallel-contact bipolar stimulating electrode was implanted at interaural coordinates AP +3.4 to +3.6; ML +1.6; DV 2.1 to 2.3. Substantia nigra stimulation consisted of 50 biphasic pulses (0.5 ms duration) applied at 100Hz (average current applied for each group 500 to 990 uA).

### Visual stimulation

In the extracellular recording experiments, visual stimuli (10 ms duration, 0.2 Hz) were delivered by a white LED (1500 mcd) that was placed 1-2 cm directly in front of the right eye of the animal. The left eye was covered. LED and electrical stimulating electrodes were connected to constant current electrical stimulators (Isolator-10, Axon Instruments Inc.).

### Bicuculline injections

The drug-filled pipettes were lowered to 4.0-4.2 mm from brain surface into the deep layers of the superior colliculus (AP −6.5/ ML +1.5 mm), and either supported by the IVM micromanipulator or secured with dental cement. Bicuculline (0.01% in saline, 250 nl) was injected into the superior colliculus at a rate of 400 nl/min.

### Extracellular recording

Extracellular single unit recordings were made using 5-15 MΩ micropipettes. Electrodes were filled with 1M NaCl solution with 2% neurobiotin (SP1120, Vector). Only stable neurons with wide average spike wave form (>1.1ms), and slow spontaneous firing rate (<0.1 Hz) were included. Neurons with train spike activity typical of low threshold-spiking (LTS) neurons were also excluded from this study. The spike rate of the recorded SPNs during 30 ms following each cortical stimulation was used as an indication of the strength of corticostriatal synapses. Recordings were made via either a headstage (model HS-2A) connected to an Axoprobe-1A microelectrode amplifier (Axon Instruments Inc California,

USA), or a headstage (NL 100 Neurolog) connected to a preamp (NL104), an amplifier (NL106) and a filter (NL125). Signals were amplified and band-pass filtered (0.1 to 10,000 Hz). All waveform data were digitized at 50 kHz by an A-D interface (1401 Micro 2, CED, UK), and acquired using SPIKE2 software (v6 or v7, CED).

### Intracellular recording

Intracellular recordings were made using 35 to 130 MΩ micropipettes with 1 M K-acetate internal solution, in some cases containing 3 to 4% biocytin. For intracellular recording, only stable neurons with a membrane potential more negative than –60 mV that displayed characteristic spontaneous fluctuations in membrane potential (> 10 mV amplitude) and action potential firing were included in this study. Current-voltage relations were obtained by injecting hyperpolarizing and depolarizing current pulses through the micropipette, using an Axoclamp-2B amplifier (Molecular Devices) configured in current-clamp mode. Membrane potential fluctuations were recorded for periods of at least a minute after the cell had stabilized following impalement and at regular intervals of >15 min. All waveform data were digitized at 10 kHz by a Digidata 1200B or a Digidata 1322A (Molecular Devices), displayed using pClamp 8 software (Molecular Devices).

### Extracellular recording experimental protocol

After a stable single unit recording was obtained from a putative spiny projection neuron, cortical stimulation (0.2 Hz) was applied throughout the recording, and the short latency (<30 ms) spike response was measured as the strength of the corticostriatal synaptic input. During the recording, firstly, a baseline of 10 minutes of spike activity was recorded. The cortical stimulation was then paired with light stimulation for another 5 min. Bicuculline was then injected locally to the deep layers of the superior colliculus. Visual stimulation was paired for another 10 min with the cortical stimulation. In the match group, visual stimulation was applied when the iLFP started to decrease, approximately 500 to 600 ms after cortical stimulation. In the mis-match group, visual stimulation was applied when the iLFP started to increase, approximately 250 ms after cortical stimulation. Therefore, the estimated light-induced phasic dopamine activity occurred either during the ChI pause (Match group) or excitation (Mis-match group, Figure 1C). In the control group, the visual simulation was not applied after BIC injection. After pairing, the recording of spike responses to cortical stimuli continued for at least another 30 minutes.

The peristimulus histograms (PSTH) of putative ChIs and SPN spikes were plotted using the cortical stimulation (Figure 1C, 1D) or the trough of the iLFP (Figure 2C) as the triggers. Each PSTH represents 60 sweeps of recording and the bin sizes were 50, 1, and 20 ms in Figure 1C, 1D and 2C respectively.

### Intracellular recording experimental protocol

After a stable impalement was obtained from a spiny projection neuron, a baseline of 20 minutes of postsynaptic potentials (PSPs) was recorded before the plasticity-inducing protocol. Postsynaptic potentials were always elicited from the Down state. The transitions between membrane potential “states” were detected online using a locally-constructed functional clamp^1^, and this used to trigger data acquisition and substantia nigra electrical stimulation, where relevant, early in the ensuing Down or Up state. A delay of 600 ms was imposed between intracellular current injection and substantia nigra stimulation in the dissociated group to ensure that the depolarization was induced in the up state, and the substantia nigra stimulation occurred primarily in the ensuing down state. Intracellular current injection, when included in the protocol, was set 0.2 nA higher than the threshold for eliciting continuous action potential firing (range 0.5 to 2.0 nA). Membrane potential fluctuations and current-voltage relations were assessed before and after the plasticity-inducing protocol to ensure that changes in PSPs were not associated with changes in membrane characteristics. The experimental protocols used are illustrated in Fig 2C.

### Histology

At the end of extracellular recording, the putative spiny neurons was actively filled with neurobiotin by a juxtacellular filling protocol^2^. Briefly, the target cells were “driven” to spike by applying positive current through the recording pipette (up to 12 nA, 250ms on-off pulses) for up to 15 min. At the end of intracellular recording, putative spiny neurons were filled intracellularly with biocytin by applying depolarizing current pulses (0.8 to 1.5 nA; 3 Hz; 10 to 15 min) via the recording micropipette.

Vibratome sections (50 - 60 µm) were processed using standard histological procedures ^3^ and labeled cells were identified by light or fluorescent microscopy. In the extracellular recording experiments, 8 neurons, and in the intracellular recording experiments, 14 neurons, were recovered and verified histologically as spiny projection neurons.

Positions of the substantia nigra electrodes for intracellular recordings were determined in unstained or cresyl-violet stained sections to be within 500 µm of the dopamine cell layer of the substantia nigra pars compacta. There were no systematic differences in electrode positions between groups. In groups where SN stimulation was not applied, electrodes were still fitted during surgery to control for the release of dopamine that may accompany acute electrode implantation^4^.

### Data analysis

Data were analyzed offline using Axograph 4.9 software and SPIKE2 v6 or v7. Statistical tests on data from single cells as well as on group data were performed in SPSS and Prism.

